# sPreserved recognition of basic visual features despite lack of awareness of shape: evidence from a case of neglect

**DOI:** 10.1101/2023.05.30.541805

**Authors:** Seda Karakose-Akbiyik, Teresa Schubert, Alfonso Caramazza

**Affiliations:** Department of Psychology, Harvard University, Cambridge, Massachusetts, USA; Center for Mind/Brain Sciences – CIMeC, University of Trento, Rovereto, Italy

## Abstract

Human visual experience of objects comprises a combination of different visual features, such as their color, position, and shape. Spatial attention is thought to play a role in creating a coherent perceptual experience, integrating visual information coming from a given location, but the mechanisms underlying this process are not entirely understood. Deficits of spatial attention in which this integration process does not occur normally, such as neglect, can provide insights regarding the mechanisms of spatial attention in visual object recognition. In this study, we describe a series of experiments conducted with an individual with neglect, XX. XX presents characteristic lack of awareness of the left side of individual objects, evidenced by poor object and face recognition, and impaired word reading. However, he exhibits intact recognition of color within the boundaries of the same objects he fails to recognize. Furthermore, he can also report the orientation and location of a colored region on the neglected left side despite lack of awareness of the shape of the region. To our knowledge, selective lack of awareness of shape despite intact processing of basic visual features in the same spatial location has not been reported previously. XX’s performance raises intriguing questions and challenges about the role of spatial attention in the formation of coherent object percepts and visual awareness.

## Introduction

The visual system is bombarded by an immense amount of information at every glance. Decades of research have revealed that dedicated networks extract and process information such as orientation, color, and motion of stimuli at every location in the visual field to create a coherent visual experience (Hubel & Wiesel, 1968; Livingstone & Hubel, 1988; Zeki et al., 1991). The visual system achieves this feat in part by segmenting and processing individual objects and processing their various properties in parallel (e.g., a mug, red, blue stripes). Spatial attention is thought to play a role in creating a coherent perceptual experience (a red mug with blue stripes) by integrating visual information coming from a given location (Humphreys et al., 2000; Kahneman et al., 1992; Robertson, 2003; Treisman & Gelade, 1980; Wolfe & Bennett, 1997; Wolfe & Horowitz, 2004). Despite this insight, the mechanisms underlying this integration process are not entirely understood.

One source of insight regarding the mechanisms of spatial attention in visual recognition comes from deficits of spatial attention, in which the integration of object features does not occur normally. In this study, we examined the perception of different visual features (shape, color, and orientation) in an individual with neglect, XX. Through study of his patterns of behavior, we gain insight into how visual features of objects are accessed and combined in visual object recognition, and the mechanisms of spatial attention in this process.

Neglect is an acquired disorder of spatial attention, often resulting from damage to right hemisphere parietal and/or frontal cortices (Bisiach, 1993; De Renzi, 1982; Vallar & Perani, 1987). In the absence of sensory processing deficits, individuals with neglect typically lose awareness of visual information on the side of space opposite to their lesion, the contralesional side (for reviews, see Lunven & Bartolomeo, 2017; Vallar & Calzolari, 2018). For instance, an individual with a right hemisphere lesion might ignore the left side of an object (e.g., fail to notice and eat food on the left side of their plate) or make errors in reading on the left side of a word (e.g., read *dice* as “*race*”).

The statement that neglect affects the *left half of an object* already implies that there must be sufficient processing of the entire object so that its boundaries and center can be defined by the visual system. Supporting this idea, individuals with neglect can segment images based on low-level visual information, including information present on the neglected side (Driver & Mattingley, 1998). As an example, when asked to place a mark in the middle of a line, individuals with left neglect tend to place their mark rightward of the center (Bisiach et al., 1983), as expected based on lack of awareness of the left half of the line. However, the amount of rightward bias increases proportionately to the length of the line, reflecting some degree of visual processing of the entire line to scale accordingly (Ro & Rafal, 1996; Vallar et al., 2000; for similar evidence in word reading see Caramazza & Hillis, 1990; Kinsbourne & Warrington, 1962; Schubert & McCloskey, 2013; Subbiah & Caramazza, 2000; Young et al., 1991). Overall, there is evidence that basic perceptual processes such as figure-ground segmentation can be carried out without awareness and this segmentation can influence task responses (for review of similar evidence in neurotypical subjects, see Vandenbroucke et al., 2014).

In addition to implicit processing of spatial extent, individuals with neglect can also show implicit knowledge about the nature of stimuli or stimulus parts they fail to report, as documented with indirect measures such as priming. For instance, individuals with neglect might fail to detect an object on the left side of the visual field (e.g., a drawing of an apple), yet respond faster to a subsequently presented word if it is semantically related to the missed object (e.g., after apple, faster for *tree* compared to *truck*, Audet et al., 1991; Berti & Rizzolatti, 1992; Kristjánsson et al., 2005). Thus, there is also evidence that neglected stimuli can be processed implicitly to some extent and affect later processing of other objects.

Taken together, previous research into information processing in neglect has revealed a dissociation between the availability of implicit and explicit information about visual stimuli on the neglected side. The evidence for figure-ground segmentation and implicit encoding of stimulus properties on the left side shows that neglected visual properties can be unconsciously processed to a certain extent regardless of whether they are consciously perceived. This pattern supports the long-held view that focused spatial attention is required for overt recognition of stimulus properties (Beck et al., 2001; Deouell, 2002; Driver & Mattingley, 1998; Driver & Vuilleumier, 2001; Rafal, 1994; Treccani et al., 2012; Van Vleet & Robertson, 2009; Vuilleumier et al., 2001). In previous work, it has been assumed that when an individual with neglect is unaware of visual information at a given location, they are experiencing a failure to attend that prevents overt recognition of *all information* residing in that location.

In this study, we report a case of left-neglect, an individual we refer to as XX, who shows selective lack of awareness of left-side information about object shape while retaining the ability to report color defined within the same object. Furthermore, he can also report the location and orientation of a colored region on the neglected left side despite lack of awareness of the shape of the region. To our knowledge, selective impairment for some but not all features in the same spatial location has not been reported in other cases of neglect. XX’s behavioral profile thus raises new questions regarding how visual features of objects are accessed in visual object recognition, and the role of spatial attention in this process.

## Results

### A case of left neglect: XX

XX is a right-handed man who was 67 years old at the start of data collection in May 2019. He has some college education (no degree) and does not report any premorbid cognitive disabilities. In 1988, he suffered a cerebral hemorrhage that resulted in complete left hemiparesis and problems in visuospatial processing. His symptoms suggest a lesion to the right parietal lobe, including involvement of the motor strip; a complete description of his brain lesion is not available, and neuroimaging could not be obtained due to metal implanted during surgery.

Our neuropsychological evaluation did not reveal any difficulties with his semantic or linguistic processing, verbal short-term memory, or basic visual perception (see SI Appendix A). Screening of neglect was carried out using standard screening batteries (unlimited duration stimulus presentation) as well as custom experiments (limited duration stimulus presentation). Despite relatively good performance on some neglect screening tests with unlimited duration, XX’s performance was clearly abnormal and showed signs of neglect on the left side (see SI Appendix B). For instance, in direct copying of a single flower, he missed the petals on the left (see Figure S1). In visual search tasks, he started searching for items on the right and moved to the left only after completing the right side. The main experiments for this study were custom computerized tasks with limited exposure durations, which allowed us to gain a more detailed understanding of XX’s deficits (see Methods for details).

### XX neglects the left side of objects, faces, and words

As an initial test of the extent and nature of neglect in object perception, we used black and white line drawings of objects or chimeric figures that combine right and left halves of different objects (Experiment 1a, see Figure 1A). XX’s task was to indicate whether the figures were real or made-up objects. To perform accurately in this task, both sides of the figure must be processed to determine if they differ (Buxbaum & Coslett, 1994; Hillis & Caramazza, 1995; Subbiah & Caramazza, 2000; Young et al., 1992).

**Figure 1.**
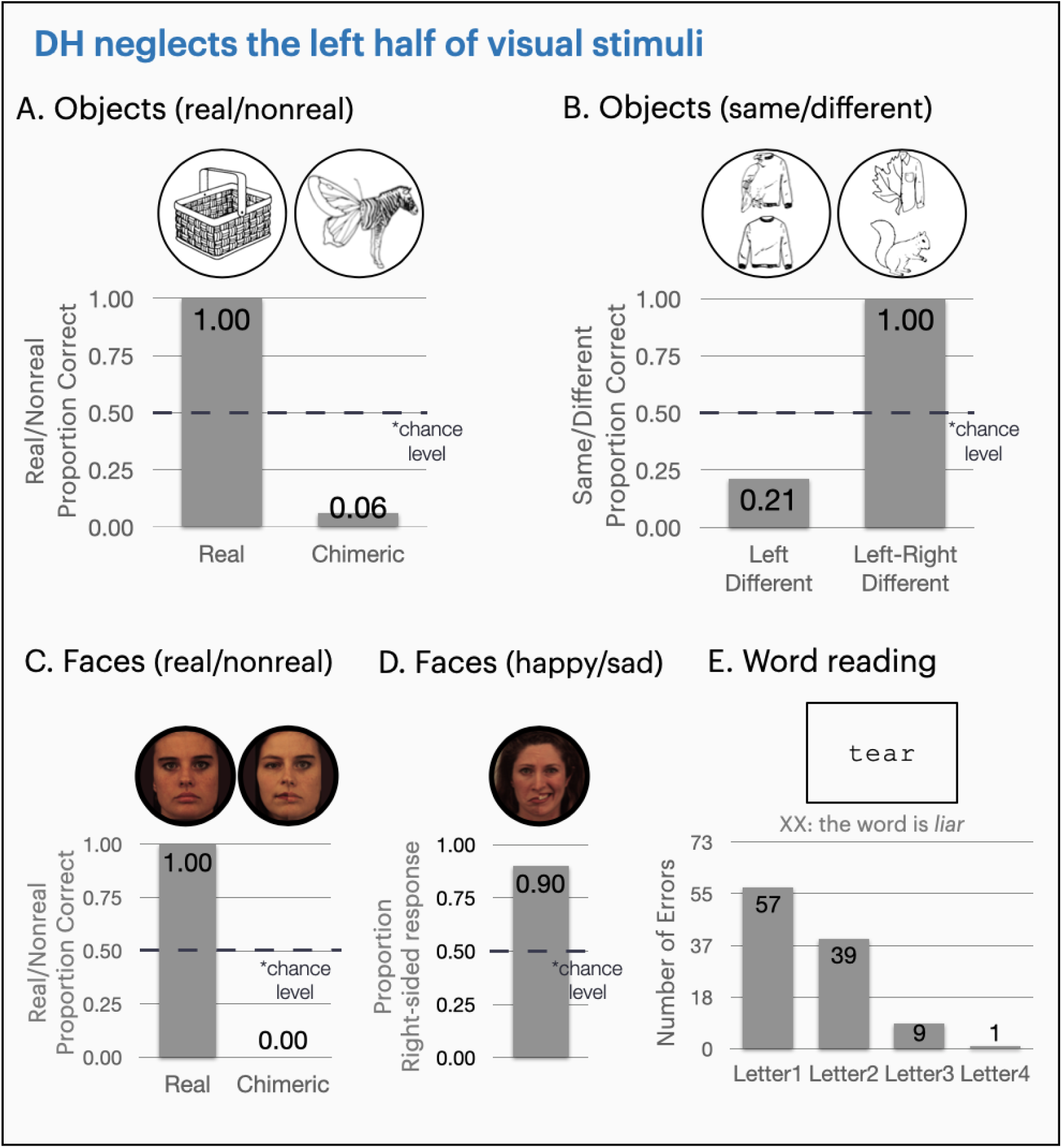
Summary of XX’s performance in tasks that require perceiving the left half of objects, faces, and words. **(A)** Object decision and **(B)** same-different task performance for black and white line drawings of objects or chimeric figures. Both tasks revealed lack of awareness of shape information on the left. Face perception tested by **(C)** chimeric faces and **(D)** chimeric facial expressions. XX does not notice the aberrant left sides and responds solely on the right sides of face stimuli. **(E)** Distribution of letter substitution errors by position in word reading. For ease of comparison across different letter positions, only the errors that preserve word length are displayed in this graph (82% of errors). Note that he tends to make letter substitution errors on the left half of words (e.g., reads the word *tear* as “*liar*”, or *race* as “*dice*”).

When presented with these stimuli, XX spontaneously named all real objects and identified them as real (100% correct), but also misidentified all except one chimeric figure as real, naming them based on the object depicted in the right half (see Figure 1A, real/nonreal task: 6% correct for chimeric figures, below chance, *p* < .001). For instance, for the chimeric figure shown in Figure 1A, he responded “*this is a real zebra*”. As another demonstration of his failure to notice the left sides of visually presented objects, we conducted a same-different task with two vertically aligned images (objects or chimeric figures) presented at the center of the screen (Experiment 1b). XX’s task was to indicate whether the two images were the same or different. He did not need to recognize the images to succeed in this task: all he needed to identify was if there was any visual difference between the two images. For these stimuli, XX answered solely based on the right sides, describing any two stimuli as identical if their right sides matched (see Figure 1B, same/different task: 100% correct when both halves were different; 94% correct for identical pairs; 21% correct for pairs that differed only on the left side, below chance, *p* = .019).

Overall, XX’s responses reflected a problem in processing shape information on the left half of visually presented objects. Next, we investigated whether this difficulty extended to other classes of objects such as faces and words. First, we created chimeric faces that combine right and left halves of two different faces and presented them along with unaltered faces (Experiment 2a). To test if he could process both halves of a face, we asked him to report whether the faces were real (see Figure 1C). XX endorsed both the chimeric and real faces as real (real/nonreal task: 100% correct for real; 0% correct for chimeric), consistently missing the mismatch between the left and right halves of the face.

Next, we used emotionally chimeric faces that combine the right and left halves of happy and sad expressions of the same person, presented along with whole emotional faces (Experiment 2b, see Figure 1D). XX’s task was to report the emotion on the faces as happy or sad. He reported the correct emotion in 97% of the whole faces. However, for chimeric faces, he reported the emotion on the right side in 90% of responses and did not notice the aberrant left sides. For instance, in response to the image in Figure 1D that is happy on the right and sad on the left side, he responded “*she’s happy”*. He was equally confident in his responses regardless of whether the face was chimeric or whole (rated on a scale of 1 to 5, *M_chimeric_* = 3.72, *SD_chimeric_* = .83; *M_real_* = 3.97, *SD_real_* = .81, *t*(76) = -1.38, *p* = .17, see Methods for more detail).

To examine XX’s performance in word reading, 4-letter words were presented (Experiment 3). In order to reduce the possibility of getting a word correct just by chance, the list included pronounceable nonwords and words with different sizes of lexical cohort (i.e., other words sharing the beginning or end of the word, e.g., *nail*’s cohorts are *rail*, *fail,* etc.). He made errors in word reading in 46% of trials, 82% of which preserved the length of the word (e.g., reading *face* as “*dice*”). Analyzing his letter substitution errors by letter position, we observed a characteristic pattern of neglect where he tended to make letter substitution errors on the left side of words but rarely on the right side (e.g., reading *easy* as “*busy*”, see Figure 1E).

XX had difficulty in perceiving visual information on the left half of objects, faces, and words in a variety of tasks with stimuli presented centrally. Due to reports of previous patients that neglect either the left half of the visual field or individual objects (Calvanio et al., 1987; Caramazza & Hillis, 1990; Medina et al., 2009; Vallar & Maravita, 2003; Verdon et al., 2010), we ran a series of experiments addressing whether his deficit is defined by the visual field or applies to individually segmented objects (*retinocentric* versus *allocentric* neglect, for more detail, see SI Appendix C, Figures S3-S6, Supplemental Methods Experiments 4a-e). These experiments revealed that XX neglects the left half of individually segmented objects, and his problem in processing the left half of objects persists even when the entire stimulus falls in the right visual field.

#### Reporting visual features on the neglected side: color

We showed that XX neglects the left half of individually segmented objects, faces, and words. Does the deficit to attend to one half of an object harm processing of all visual features, or can different visual features be accessed and reach awareness independently? To address this question, we systematically varied basic visual features on the left half of stimuli. We first tested color due to its specialized circuitry in early visual processing and because it can draw attention independently of other visual features (D’Zmura, 1991; Treisman & Souther, 1985; Wolfe & Bennett, 1997). We generated stimuli that varied in color on the left and right side (bicolor words and bicolor chimeras) and tested XX’s performance in reporting shape- and color-related information. Given XX’s deficit in attending to the left half of objects, and given previous reports of neglect patients, we would expect his responses to not reflect any information coming from the left half of an object. For instance, for a figure that is half a swan in red on the left half and half a truck in blue on the right side, we would predict that he would neglect all visual information on the left half of the image as in the previous experiments and report “*a real blue truck*” (see Figure 2A).

**Figure 2.**
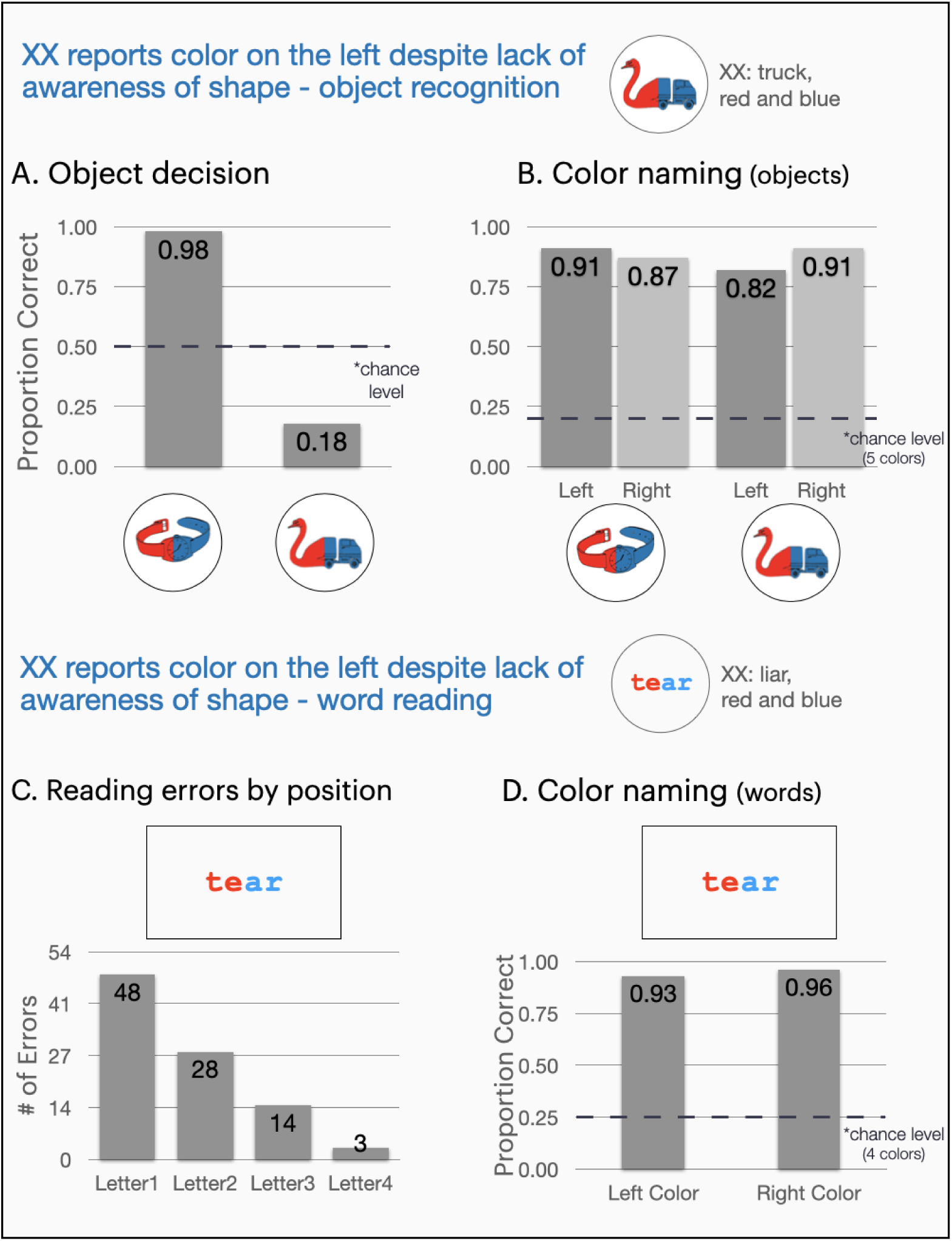
**(A)** Object decision and **(B)** color naming performance for bicolor chimeric figures. Color naming performance is presented on both left and right sides of stimuli for comparison. Despite neglecting the left halves of chimeric figures, XX can report the color on the same location. **(C)** Distribution of letter substitution errors in reading bicolor words. To ease comparison across letter positions, only errors that matched the target word length are presented (79% of errors for bicolor words). **(D)** Color naming performance on the left and right half of bicolor words. Despite left-sided letter substitution errors in word reading, XX can report the color on the neglected left side.

### Dissociating color and shape processing on the neglected side: colored chimeras

Bicolor line drawings of real objects and chimeric figures were created by coloring the left and right half of line drawings with a different color (Experiment 5, see Figures 2A, B) and presented along with unicolored figures. XX again completed an object-decision task (i.e., decide whether the line drawing is of a real object), but was required to also report the colors of stimuli. Overall, he correctly endorsed 99% of real objects but also endorsed most chimeric figures as real (real/nonreal task: 26% correct, below chance, *p* < .001). His object decision performance for chimeric figures was not significantly different across unicolor (33% correct) and bicolor conditions (18% correct, *X^2^* = 2.00, *p* = .16). Thus, the difference in color between the two halves of the bicolor chimeric figures did not help him perceive the left side of the object. Strikingly, despite his failure to perceive the aberrant left side of chimeric figures, he was able to report color information on the left half of the object (see Figure 2B, all values above chance, all *ps* < .001). Furthermore, in 81% of bicolored chimeric figures that he misidentified as real, he accurately reported the left color (sample response for the bicolor chimeric stimulus in Figure 2A: “*a real truck, red on the left, blue on the right*”). Thus, XX demonstrated preserved recognition of color in the otherwise-neglected half of objects. As we have already shown that he neglects the left half of words, we then addressed whether this dissociation between shape and color on the left also holds for words.

### Dissociating color and shape processing on the left: colored word reading

4-letter words were presented in bicolor and unicolor conditions (Experiment 6, see Figure 2C) and XX’s task was to read the words and name their colors. His word-reading performance in the two conditions was comparable (bicolor words: 41% correct, unicolor words: 49% correct, *X^2^* = .79, *p* = .37), suggesting that the difference in color between the two halves of the word did not help him perform better on the left side. He made typical left-sided letter substitution errors for both bicolor and unicolor words (e.g., reading *perk* as *bark* in the bicolor and *pork* in the unicolor condition, see Figure 2C). However, in both bicolor and unicolor conditions, he generally reported the color on the left correctly (bicolor: left 93%, right 96%; unicolor left 96%, right 99%, all above chance, *ps* < .001, see Figure 2D). Importantly, in 92% of trials in which he made an error in reading a bicolor word, he accurately reported the color on the left. For instance, in reading the word *perk* that is yellow on the left (letters *p* and *e*) and red on the right (letters *r* and *k*), he reported “*bark, yellow and red*”.

#### Testing boundary conditions of color perception on the neglected side

XX’s responses to colored chimera and words revealed a striking dissociation in processing color and shape on the left half of visually presented stimuli. Despite the characteristic difficulty in processing the left half of objects and words, XX was able to report color information on the left side. Previous work revealed a gradient of attentional performance in neglect such that attentional deficits become more pronounced as distance from the center increases (Behrmann et al., 1997; Halligan et al., 1992; Kinsbourne, 1987). Thus, for both colored chimeras and colored words, it was possible that information near the midline might have been enough for XX to recognize the color on the left even if he did not process the color of the entire figure. To address this possibility, we created stimuli that vary the size and position of the two colors in the figure (Experiments 7a-b). Since previous work showed better allocation of attention to regions of higher saliency (Fellrath & Ptak, 2015), varying the size of the colored regions also addressed if the saliency of the colored patch would affect XX’s success in color recognition (Experiment 7a). We found that XX performed at ceiling in recognizing the left-side color, even when th oddly colored portion comprised only 1/10^th^ of the entire stimulus and/or was at the far-left edge of the stimulus (see Figures 3A-B).

**Figure 3.**
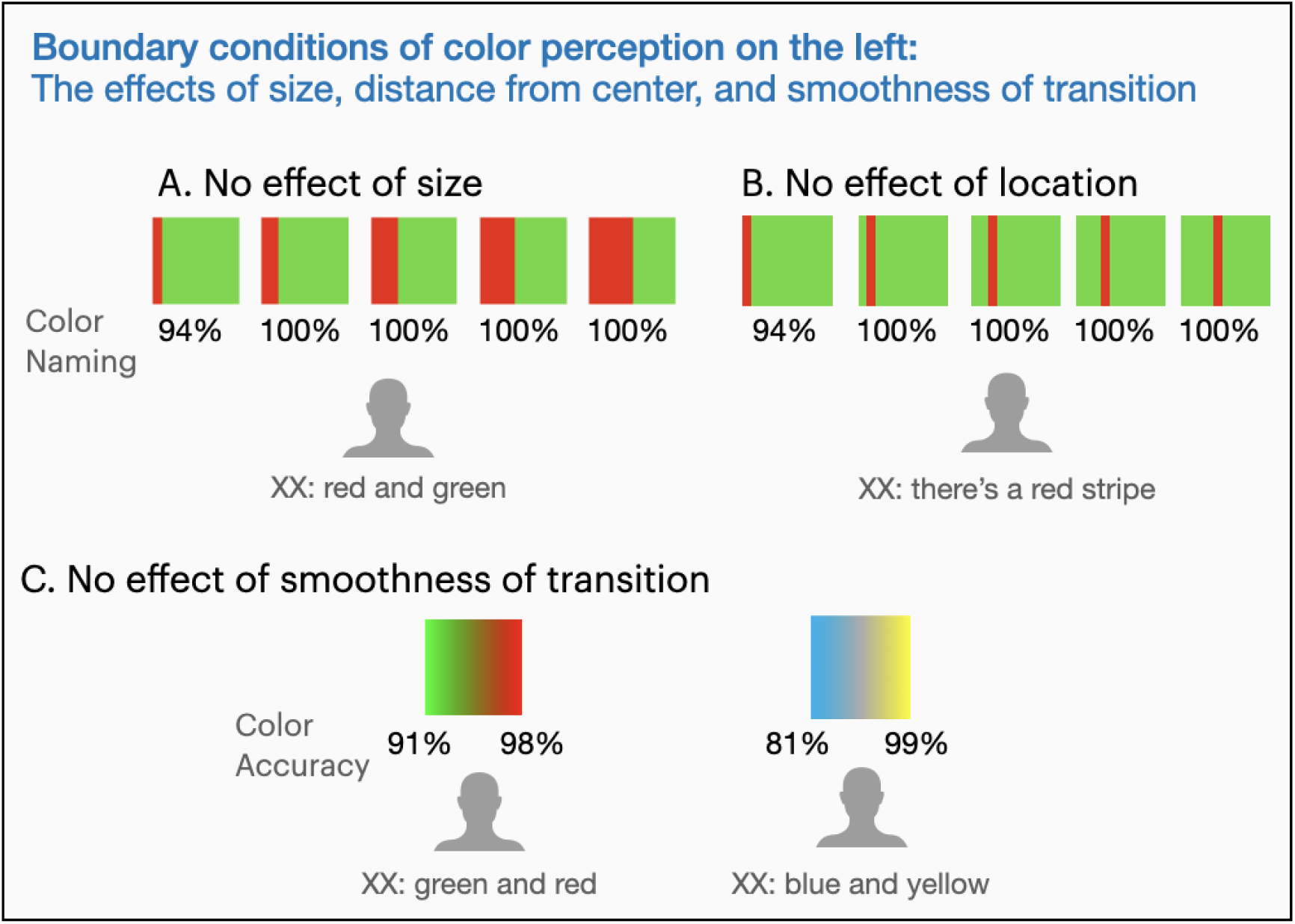
Experiments testing the boundary conditions of color perception on the left side. To test the effects of salience and distance from center, bicolor squares were used where **(A)** the two colors varied in their proportion within a stimulus, **(B)** and a colored patch of a given size changed location in 1/10 increments (1/10 of the whole square). XX performed at the ceiling in recognizing the left-color even when the oddly colored portion comprised only 1/10^th^ of the entire stimulus and was at the far-left edge of the stimulus (94%). Note that we are reporting data from 1/10^th^ to 5/10^th^ conditions to demonstrate his performance on the left; he was 100% correct in reporting the color on both sides starting from 2/10^th^ of the left side onwards. To control for the possible segmentation of the colored squares into separate-colored rectangles, **(C)** we used bicolor squares where the color transition was smooth (*left side*: the colors faded into each other, *right side*: the colors faded into gray in the middle). Here too XX was able to report the color on the neglected left half.

XX was able to report the color on the left even when the colored portion comprised a small part of the stimulus or was far from the center of the object. However, an abrupt change of color between two portions of an image could potentially result in two distinct representations for the two halves (Palmer & Rock, 1994; Vannuscorps et al., 2021). That is, XX’s visual system might have divided the bicolor shape into two individual stimuli or objects, which then could have supported the reporting of their color even if the left half of each stimulus was neglected. To address this possibility, we ran two experiments with continuous color gradients that eliminate the abrupt change in color (Experiments 7c-d, see Figure 3C). In these experiments, XX was still able to report the color on the left side. Thus, his success in reporting the color on the left cannot be explained by a possible segmentation of the colored squares into separate-colored rectangles.

#### The relationship of color to other visual features on the neglected field: shape, orientation, and location

XX can accurately report color on the neglected left side despite lack of awareness of shape information. Can he recognize other stimulus properties on the left, such as orientation, or is his success specific to color? There is evidence for specialized neural circuitry for orientation (Ferster & Miller, 2000) and that orientation can draw attention independently of other visual features (Treisman, 1986; Wolfe et al., 1992; Wolfe & Horowitz, 2004). Furthermore, encoding of orientation can dissociate from object identity or shape (Vannuscorps et al., 2021). Given these considerations, we tested orientation as a case study of contralesional visual processing of other features. To contrast XX’s recognition of color, orientation, and shape, we created stimuli that allowed a direct comparison of these features. Finally, following previous proposals on the link between the encoding of stimulus location and awareness (Allport, 1987; Berti, 2002; Berti & Rizzolatti, 1992; Rizzolatti & Gallese, 1988), we also tested whether XX can report the position of colored stimuli on the neglected side.

### Integration of color and simple geometric shapes

We presented chimeric figures or objects with a colored patch (circle or square) on the left or right side (Experiment 8, see Figure 4A). When asked to identify if the figures were real, XX correctly endorsed 100% of the real objects as real but also endorsed nearly all chimeric figures as real objects (7% correct, below chance, *p* < .001). Despite this characteristic difficulty in object decision, he was able to report the color of the patch when it appeared on either the left or right side (see Figure 4A, above chance, all *ps* < .001). Consistent with his performance in colored chimeras, in those chimeric trials where he neglected to notice the aberrant left sides of chimeric figures, he could still recognize the color of the patch presented on the left side (93% correct, above chance, *p* < .001).

**Figure 4.**
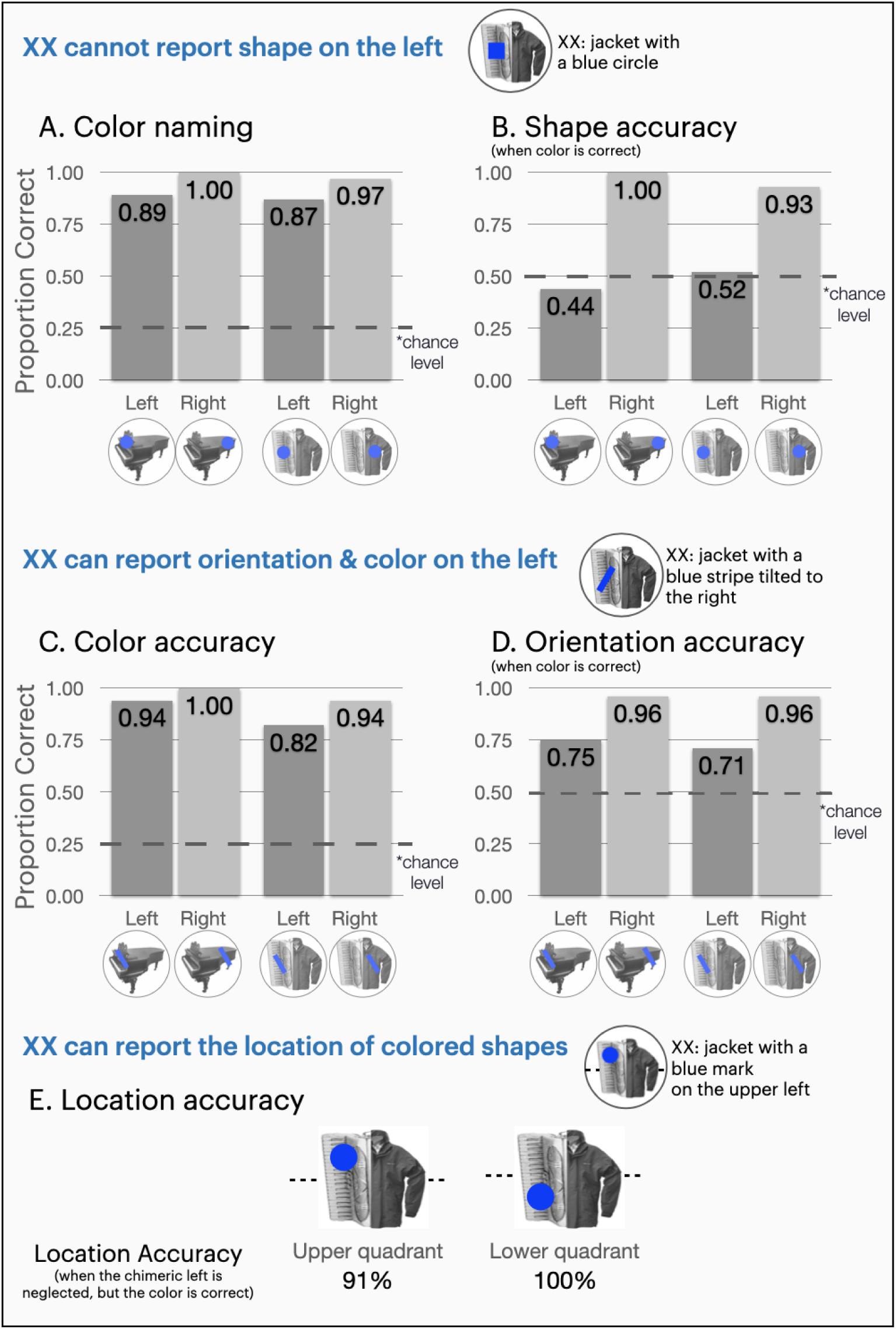
Experiments testing the integration of color with shape, orientation, and location, using real objects and chimeric figures. Color and orientation accuracy are presented on both left and right sides of figures for comparison. **(A-B)** *Integration of color and shape.* (A) Color naming performance when a colored patch was presented on real objects or chimeric figures. XX was able to report color on the left without difficulty. (B) Circle/square shape discrimination performance on real objects or chimeric figures when the color was reported accurately. Despite being able to report the color of a geometric shape on the left, XX was at chance at reporting whether the shape was a circle or a square. **(C-D)** *Integration of color and orientation.* (C) Color naming performance when a colored rectangle was presented on real objects or chimeric figures. XX could report color on both sides without difficulty. (D) Left/right orientation accuracy of a rectangle when its color was reported accurately. When XX reported the color of a rectangle on the left, he could also report its orientation. **(E)** Encoding of stimulus location on the neglected left halves of chimeric figures when the color of the stimulus was recognized. Despite neglecting the left halves of chimeric figures, XX could report the color and position of a colored shape appearing on the same side.

Importantly, for both chimeric figures and real objects, in those trials where he correctly reported the color of a patch on the left, he was at chance in saying if it was a square or a circle (real: 44%, at chance, *p* = .69; chimeric: 52%, at chance, *p* > .99, see Figure 4B). For instance, on a given trial in which a chimeric figure was presented with a blue circle on the left, he would report the figure was real and be equally likely to describe the patch as a “*blue square*” or a “*blue circle*”. This pattern is in line with his performance in colored chimeras or colored word reading where he neglected shape information on the left but could still accurately report color.

### Integration of color and orientation

To test if XX can report orientation on the left side, we presented chimeric figures or objects with a colored tilted rectangle (tilted towards left or right for 30 degrees) on the left or right side (Experiment 9, see Figure 4C-D). When asked to identify if the figures were real, XX correctly endorsed 99% of the real objects as real but also endorsed nearly all the chimeric figures as real objects (9% correct, below chance, *p* < .001). Despite this failure in object decision, he accurately reported color information on both sides (see Figure 4C, all above chance, *ps* < .001). Consistent with his performance in colored chimeras, in those chimeric trials where he failed to recognize the aberrant left half, he could still recognize the color of the rectangle presented on the same side (84% correct, above chance, *p* < .001).

However, unlike in the previous experiment, XX was able to report more than just the color of the colored patches. Here, recognition of color and orientation tended to go together: whenever XX was able to report the color of the rectangle on the left, he was able to report its orientation as well (see Figure 4D, real objects: 75% correct, above chance *p* < .001; chimeric figures: 71% correct, above chance *p* = .008). Most importantly, on those chimeric trials where he neglected the aberrant left side, if he reported the color on the left (84% of trials), he was above chance in reporting the orientation as well (75% correct, above chance, *p* = .004). For instance, for a chimeric figure comprised of the left half of an accordion and right half of a jacket with a blue left-tilted rectangle on its left, he responded “*it’s a jacket with a blue rectangle on it, the rectangle is tilted to the left*”. Overall, even though he was not able to recognize the aberrant left halves of a chimeric figure, he was able to report both color and orientation of a rectangle presented on the same neglected left side.

### Integration of color and location

Following previous proposals that encoding of stimulus location is a precondition for it to enter consciousness (Allport, 1987; Berti, 2002; Berti & Rizzolatti, 1992; Driver & Vuilleumier, 2001; Rizzolatti & Gallese, 1988; Robertson, 2003), we next addressed whether XX could report the position of colored regions on the neglected left side of an object. To this end, a colored circle was presented at four different locations (left/right, upper/lower quadrant) within chimeric figures or real objects (Experiment 10, see Figure 4E). The upper or lower quadrant was specified in relation to a dashed line behind the stimulus, and XX reported the location of the circle as above or below the dashed line. When asked to identify if the stimuli were real, XX correctly endorsed 99% of the real objects as real but also endorsed most chimeric figures as real objects (13% correct, below chance, *p* < .001). Thus, he showed the typical lack of awareness for object shape on the left side.

As for the colored circles located within the object, he failed to detect a colored patch in 24% of trials when it was presented on the left upper quadrant, and 48% of trials on the left lower quadrant. Thus, he was more likely to miss the colored patch when it resided in the lower quadrant of the object (for a discussion of neglect effects on the vertical dimension of space, see Làdavas et al., 1994; Pitzalis et al., 1997). However, whenever he detected the presence of a colored patch, he was also able to report its color (upper quadrant: 95%, lower quadrant: 92%, both above chance *p* < .001).

As for location of the colored patches located within the object, in those chimeric trials where he neglected the aberrant left side, that is, showed lack of awareness of shape information on the left side, whenever he reported the color of the patch on the left, he could also report its location (upper quadrant: 91%, lower quadrant: 100%, *p*s < .001). That is, even though he neglected shape information on the left half of the object, he could report the color and the location of a colored patch on the same neglected side.

## Discussion

In this study, we documented the behavioral profile of an individual with neglect, XX, who shows characteristic neglect of the left half of visually presented objects, faces, and words. However, despite lack of awareness of shape information on the left half of visual objects, he displays a unique pattern never before reported: he is able to report color and orientation information from the neglected left side of an object. Furthermore, he can report the color and location of colored parts within the left side of an object while not being able to recognize the object itself. To our knowledge, selective impairment for some but not all visual features in the same spatial location has not been reported in any instances of neglect or other studies of attentional failure. This unique pattern of results has implications for how visual features of objects are encoded and selected in visual object recognition.

In typical perception, when an observer becomes aware of an object, its features are already bound to one another and specific locations. Visual awareness consists of complete objects comprised of color and texture, at a particular location in the visual field. Attention is thought to play a role in creating this coherent perceptual experience, and various theories of space-based and object-based attention have been proposed to account for the integrative aspect of visual perception. For instance, according to *Feature Integration Theory*, focused attention to an object’s location binds its various features into a consciously reportable unified percept (Treisman, 2006; Treisman & Gelade, 1980). Object-based theories of attention, on the other hand, account for this integration by emphasizing the *object* as the unit of attentional selection. For instance, the incremental grouping (Roelfsema, 2006) and integrated competition models (Desimone & Duncan, 1995) of object-based attention suggest that directing attention to a particular feature of an object leads to selection of its other constituent features as well. Binding of features of the attended object is thus possible and the experience of a unified percept arises. Overall, it has been generally assumed that without attention, the unified percept of an object cannot arise, and its constituent features can only be processed unconsciously but will not reach the level of conscious report.

The importance of attention for visual perception has been widely established following evidence from individuals with neglect or extinction (Deouell, 2002; Driver & Mattingley, 1998; Esterman et al., 2002; Làdavas et al., 1993; McGlinchey-Berroth, 1997; Treccani et al., 2012) and confirmed in studies with neurotypical subjects through phenomena such as *inattentional blindness* and the *attentional blink* (Hutchinson, 2019; Mack & Rock, 2000; Rensink et al., 1997). However, these previous accounts of the role of attention in visual awareness cannot fully explain XX’s behavior because they predict that attention and awareness are all or nothing: one is either aware of all visual features at a given location, or unaware. That is, these accounts propose a unitary representation of an object’s features at the level of awareness. Thus, if a part of an object is not in awareness, no visual features of that part can be reported. To explain XX’s pattern of reporting some but not all visual features at a location, we need a more nuanced account of the role of spatial attention in visual awareness, allowing the selection of shape and non-shape features of an object to dissociate even when there is attention to a specific location.

Our results suggest that attention may not be unitary: attention can be allocated to a feature at a given location, but doing so does not guarantee that all visual features present in that location will be integrated and reach awareness as part of a complete object. XX’s success in describing the location of colored patches on the left side of an object that he was otherwise unaware of provides clear evidence for this conclusion. He was able to successfully encode the color itself and its spatial location. However, he still failed to report shape information from the exact same location.

This pattern furthermore invites an interpretation that attention can independently select shape and non-shape features of an object. Differences in visual complexity of the features does not seem to explain this pattern. Take XX’s performance in identifying the color and shape of a square/circle on the left half of an object (Experiment 8, see Figure 4A). Even though he was able to report the color, he was at chance in deciding whether the colored patch was a circle or a square (i.e., he was equally likely to describe a blue square as a “*blue square”* or a “*blue circle”*). To succeed in differentiating a square from a circle, all one needs to identify is whether there are any corners or straight segments, an incredibly basic visual distinction. Similarly, while deciding whether two vertically presented images are the same (see Figure 1B), all one needs to identify is whether there is a global difference in shape at any location of the images. Despite performing at chance in these basic determinations about shape, XX was able to make precise judgments about color, orientation, and location on the neglected left side.

Turning to neural mechanisms, that could underlie a dissociation in selecting shape and non-shape features of an object, it is well established that different cortical areas of the ventral visual stream show selective processing of various stimulus properties, including color, orientation, and shape (Cowey, 1985; Grill-Spector et al., 1998; Hubel & Wiesel, 1962; Kourtzi & Kanwisher, 2001; Zeki, 1993). Various dorsal stream regions (particularly posterior parietal cortex) are thought to be crucial for overt recognition of these visual features encoded in the ventral visual stream (Beck et al., 2001; Brown et al., 2019; Buxbaum, 2006; Crick & Koch, 1998; Driver & Vuilleumier, 2001; Rees, 2001; Rees et al., 2002; Thiebaut de Schotten et al., 2005; Vallar & Perani, 1987; Vuilleumier et al., 2001). Although still debated, the role of these frontoparietal regions might be to focus attention (Bisley & Goldberg, 2003; Corbetta & Shulman, 2011; Petrides & Pandya, 2002), spatial encoding (Deouell, 2002), feature binding (Cohen & Rafal, 1991; Friedman-Hill et al., 1995), and/or process and maintain task-relevant perceptual information (Bettencourt & Xu, 2016; Xu, 2017, 2018). Following the hypothesis that the interaction of dorsal and ventral stream underlies overt recognition of visual features, if functionally specific parts of the ventral stream (e.g., those that selectively support color or shape processing) have distinct connections to dorsal stream regions associated with neglect, selective disruption to these connections could give rise to preserved awareness of color but not shape.

Overall, XX’s behavioral profile suggests that attention can selectively enable or impair access to different features co-located in space. Attentional allocation to a particular location does not ensure the integration of all the visual features present at that location. Instead, attention can independently select shape and non-shape features of an object, without necessarily integrating them into a unified percept. Thus, successful integration of surface features with a specific shape does not appear to be a prerequisite for conscious access to those features.

At this juncture, we would like to emphasize that XX has a chronic impairment and has lived with neglect symptoms for more than thirty years. Symptoms of neglect can partially improve, both during the acute phase of the disorder (within 6-12 months of brain injury) and during chronic stages (Berti et al., 2015; Corbetta et al., 2005; Farne et al., 2004). Thus, we cannot rule out that the unique pattern of XX’s performance relative to other individuals with neglect might have occurred due to recovery. However, even if this pattern only arose after some recovery, the pattern of behavior remains to be explained: There must be an underlying object recognition and attentional system that can be damaged in such a way to allow shape information to be neglected while color and other features are not. The theoretical implications of XX’s behavioral profile for cognitive processes underlying visual object recognition are not altered by potential recovery because that recovery must have taken place within the organizational principles of the cognitive system. Even if intact color recognition on the left side emerged because of recovery, a similar recovery did not occur for shape processing, a situation which further supports some degree of independent processing. If one had a goal of making inferences regarding *neglect as a syndrome*, data from XX may not be ideal to do so, but our aims are to make inferences regarding the typical cognitive system.

In conclusion, we documented an individual with neglect who can accurately report color and orientation from the left half of visually presented stimuli despite characteristic difficulty in reporting shape, evidenced by poor object recognition, face processing, and word reading. Through a series of experiments, we concluded that attention can be allocated to a feature at a given location, but this does not guarantee that all visual features residing there will be integrated and reach awareness. We speculate that selective disruptions in connections of functionally specific components of the ventral stream to dorsal stream regions implicated in spatial attention could give rise to these behavioral patterns. Future studies on attentional failures will test these hypotheses and help explain further how coherent visual percepts are formed and reach awareness.

## Methods

### Participants

XX was 67 years old at the start of data collection in May 2019. He is a right-handed man with some college education. He suffered a cerebral hemorrhage in 1988 that resulted in complete left hemiparesis and problems in visuospatial processing. His symptoms suggest a lesion to the right parietal lobe, including involvement of the motor strip; a complete description of his brain lesion is not possible due to metal implanted during surgery and the difficulty in obtaining his CT scans. The procedures reported here were approved by the Institutional Review Board of Harvard University and XX provided written informed consent.

### Evaluation of hemispatial neglect: Custom experiments

Unless noted otherwise, computerized tasks were carried out with the examiner sitting next to XX. A 13-inch laptop with Intel Iris Plus Graphics 655 was placed on a table in front of XX, centered at the midline of his body. In all tasks reported, unless otherwise noted, XX was asked to fixate on a central cross and give spoken responses to stimuli that were presented centrally for 500 milliseconds (see SI Figure S2). The relatively short exposure of the target stimuli was used to minimize the possibility of saccadic eye movements. Before the target stimulus, a prompt (a black square or star) appeared at fixation and XX had to report the prompt identity before reporting the target stimulus. Those trials in which he accurately reported the prompt were analyzed as valid trials. All other trials were excluded from the analyses since they indicate distraction from the task or a saccade right before the presentation of stimuli. For all tasks, stimuli were counterbalanced and randomly presented over the course of stimulus presentation. Binomial tests were used to compare XX’s performance against chance level dependent on the number of experimental conditions (e.g., real/nonreal, same/different, color naming).

#### Neglect in object processing

##### Experiment 1a

Black and white line drawings of objects or chimeric figures that combine right and left halves of different objects were presented against a white background (see Figure 1A). Chimeric figures were composed by joining halves of pictures drawn from Snodgrass & Vanderwart (1980). Twenty unique stimuli (10 real, 10 chimeric) and their reflections on the vertical axis were presented at fixation (40 trials in total). The stimuli differed in size while remaining within a bounding box of 480 by 550 pixels. XX’s task was to report whether the stimuli were real or nonreal (valid trials: 20/20 for real figures, 18/20 for chimeric figures).

##### Experiment 1b

Two objects or chimeric figures were presented vertically aligned at the center of the screen for 1000 milliseconds and presented against a white background (see Figure 1B, 60 stimuli in total: 20 identical pairs, 20 both-halves-different pairs, 20 left-halves-different pairs). Similar to Experiment 1a, chimeric figures were composed by joining halves of pictures drawn from Snodgrass & Vanderwart (Snodgrass & Vanderwart, 1980). The individual stimuli differed in size while remaining within a bounding box of 280 by 340 pixels. XX was instructed to report whether the two images were same or different (valid trials: 18/20 for identical pairs, 19/20 for both-halves-different pairs, 19/20 for left-halves-different pairs).

#### Neglect in face perception

##### Experiment 2a

Photographs of faces with a neutral expression from the Karolinska Directed Emotional Faces (KDEF) set (Lundqvist et al., 1998) were used to generate the real and chimeric face stimuli (see Figure 1C). For chimeric faces, the right and left halves of two individuals of the same gender were combined. The stimuli were cropped so that the differences in external features of the two halves of the chimeric faces (e.g., hair) were not visible. The resulting stimuli (349×261 pixels) were presented against a white background. A total of 40 trials were presented (10 real, 10 chimeric, and their reflections). XX’s task was to report whether the faces were real (valid trials: 20/20 for real faces, 17/20 for chimeric faces).

##### Experiment 2b

Chimeric emotional faces (535 x 471 pixels) were generated by combining the right and left halves of happy and sad expressions of the same person from the KDEF set and presented against a white background (see Figure 1D). A total of 80 trials were presented (40 chimeric: 20 happy-sad, 20 sad-happy; 40 whole faces: 20 happy, 20 sad). XX’s task was to report whether the faces were happy or sad (valid trials: 39/40 for whole faces, 39/40 for chimeric faces). Since his responses only reflected the emotion on the right side of a chimeric face (see Results), we wanted to test if he processed the aberrant left sides at some level. To this end, we asked him to rate his confidence in his response on a scale of 1 to 5 after he reported the emotion (5 = highly confident).

#### Neglect in word reading

##### Experiment 3

To examine XX’s performance in word reading, a total of 210 4-letter words were presented across different sessions. The words were presented in a 50-point size, set in lowercase Courier font and displayed in black against a white background. The list included pronounceable nonwords and words with different sizes of lexical cohort (i.e., other words sharing the beginning or end of the word, e.g., *nail*’s cohorts are *rail*, *fail,* etc.). For erroneous responses that preserved the word length, see Figure 1E for the distribution of errors as a function of letter position. Erroneous responses that did not preserve the word length generally took the form of adding letters to the beginning of the word (e.g., reading *hind* as *blind, germ* as *swarm*).

##### Experiments 4a-e

To determine whether XX’s neglect manifests at a retinocentric or allocentric reference frame, we conducted a series of experiments manipulating stimulus location and orientation. For a detailed description of these experiments, see SI Appendix C and Supplemental Methods.

#### Reporting simple visual features on the neglected side: color

##### Experiment 5 - Dissociating color and shape processing on the left: colored chimeras

Bicolor line drawings of real objects and chimeric figures were created by coloring the left and right half of line drawings in Experiment 1a with a different color (see Figure 2A). The stimuli were presented against a white background. Five colors were used (red, green, yellow, blue, pink). We presented XX with 208 trials in total (104 chimeric, 104 real figures, 52 of each bicolor). He again completed an object-decision task (i.e., decide whether the line drawing is of a real object), but was required to also report the colors of stimuli (valid trials: 91/104 for bicolor, 97/104 for unicolor stimuli). The stimuli differed in size while remaining within a bounding box of 480 by 550 pixels.

##### Experiment 6 - Dissociating color and shape processing on the left: colored word reading

A list of 112 4-letter words was presented in bicolor or unicolor conditions (224 trials in total, see Figure 2B). Four basic colors were used (red, green, blue, yellow). XX’s task was to read the words and name their colors (valid trials: 101/112 for unicolor words, 104/112 for bicolor words). The words were presented in a 50-point size, set in lowercase Courier font against a gray background.

#### Testing boundary conditions of color perception on the neglected side

##### Experiment 7a -Examining the effect of size on color recognition on the neglected left side

To determine whether there was a threshold width for color on the left side that allowed XX to report it, bicolor 200×200 pixel squares were generated where the size of the left-side portion varied in 1/10 increments (200×20, 200×40, 200×60 etc. pixel rectangles), resulting in nine unique bicolor conditions (see Figure 3A). The color combinations were generated by using four colors (red, green, blue, yellow) and the stimuli were presented against a gray background. A total of 220 trials were presented consisting of bicolor and solid color squares (nine bicolor stimuli presented 20 times each, each unicolor stimulus presented 40 times). XX was asked to report the colors on the squares (211/220 valid trials). In Figure 3A, we report data from 1/10^th^ to 5/10^th^ conditions to demonstrate his performance on the left; he was 100% correct in reporting the color on both sides starting from 2/10^th^ of the left side onwards.

##### Experiment 7b - Examining the effect of distance from center on color recognition on the neglected left side

Bicolor 200×200 pixel squares were generated for which a 200×20 pixel stripe appeared at 10 evenly distributed horizontal locations (see Figure 3B). A total of 220 trials were presented (nine bicolor stimuli presented 20 times each and 40 unicolor squares, 209/220 valid trials). The color pairs were generated by using four colors (red, green, blue, yellow) and the stimuli were presented against a gray background.

##### Experiment 7c-d - Examining the effect of smoothness of color transition on color recognition on the neglected left side

To rule out the possibility that an abrupt color transition enabled him to see color on both sides, we conducted two tasks where bicolor 200×200 pixel squares were presented against a white ground. The two colors were either (1) fading into each other in the center, or (2) gradually fading into one another by fading into gray in the center (see Figure 3C). Ninety-six trials were presented in each of these experiments and the stimuli were presented against a gray background (Experiment 7c: 90/96 valid trials, Experiment 7d: 91/96 valid trials). Four colors were used (red, green, blue, yellow), XX’s task was to name the colors within the squares.

#### The relationship of color to other visual features on the neglected field: location, orientation, shape

Experiments 8-10 were conducted online due to disruptions related to COVID-19. A 13-inch laptop with Intel Integrated UHD Graphics was used. The same trial structure in SI Figure S2 was followed. The experimenter controlled the flow of the tasks remotely and stayed in contact with XX over Zoom. The only difference between remote and in-person testing was that the experimenter was not sitting next to XX during remote testing. Before conducting new online experiments, we validated our online procedure by replicating the findings we had established in person.

##### Experiment 8 - Integration of color and simple geometric shapes on the neglected left side

Chimeric figures or objects with a colored patch (square: 97×97 pixels, circle: 97 pixels diameter) on the left or right were presented at fixation against a white background (see Figure 4A, 128 trials in total, 64 real, 64 chimeric). The patches came in four different colors (red, green, blue, yellow). XX’s task was to decide if the objects were real or chimeric and report the color and shape of the patch (valid trials: 59/64 for left patches, 53/64 for right patches).

##### Experiment 9 - Integration of color and orientation on the neglected left side

Chimeric figures or objects with a colored rectangle (width: 40 pixels, height: 160 pixels), tilted towards left or right by 30 degrees) were presented on the left or right half of the stimulus (see Figure 4B, 224 trials in total, 112 chimeric). The rectangle came in four different colors (red, green, blue, yellow). XX’s task was to decide if the objects were real or chimeric and report the color and tilt of the rectangle (valid trials: 102/112 left side, 106/112 right side).

##### Experiment 10 - Integration of color and location on the neglected left side

To address whether XX can report the position of colored stimuli on the left side, a colored circle (35 pixels radius) was presented at four different locations (left/right, upper/lower quadrant) of chimeric figures or real objects (see Figure 4C, 320 trials in total, 160 real, 160 chimeric). The patches came in four different colors (red, green, blue, yellow). The upper or lower quadrant was specified in relation to a dashed line behind the stimulus. XX’s task was to indicate whether the chimeric figure is real and report the color and location of the circle as above or below the dashed line (valid trials: 150/160 left side, 154/160 right side).

## Acknowledgments

We express our sincere gratitude D.H. for taking part in this study. His dedication, patience, and amazing humor made each session a truly meaningful and anticipated experience. We also thank Joseph Degutis for referring D.H. to us, and to the members of the Cognitive Neuropsychology Lab at Harvard University for their invaluable insights and feedback.

## References

Allport, D. A. (1987). selection for action: some behavioral and neurophysiological considerations of attention and action, in perspectives on perception and action, AF, HHaS, Editor. Heuer H., Sanders AF.

Audet, T., Bub, D., & Lecours, A. R. (1991). Visual neglect and left-sided context effects. Brain and Cognition, 16(1), 11–28.

Beck, D. M., Rees, G., Frith, C. D., & Lavie, N. (2001). Neural correlates of change detection and change blindness. Nature Neuroscience, 4(6), 645–650.

Behrmann, M., Watt, S., Black, S. E., & Barton, J. J. (1997). Impaired visual search in patients with unilateral neglect: an oculographic analysis. Neuropsychologia, 35(11), 1445–1458.

Berti, A. (2002). 6.1 Unconscious processing in neglect. The Cognitive and Neural Bases of Spatial Neglect, 313.

Berti, A., Garbarini, F., & Neppi-Modona, M. (2015). Disorders of Higher Cortical Function. In Neurobiology of Brain Disorders (pp. 525–541). https://doi.org/10.1016/b978-0-12-398270-4.00032-x

Berti, A., & Rizzolatti, G. (1992). Visual Processing without Awareness: Evidence from Unilateral Neglect. Journal of Cognitive Neuroscience, 4(4), 345–351.

Bettencourt, K. C., & Xu, Y. (2016). Understanding location- and feature-based processing along the human intraparietal sulcus. Journal of Neurophysiology, 116(3), 1488–1497.

Bisiach, E. (1993). Mental representation in unilateral neglect and related disorders: the twentieth Bartlett Memorial Lecture. The Quarterly Journal of Experimental Psychology. A, Human Experimental Psychology, 46(3), 435–461.

Bisiach, E., Bulgarelli, C., Sterzi, R., & Vallar, G. (1983). Line bisection and cognitive plasticity of unilateral neglect of space. Brain and Cognition, 2(1), 32–38.

Bisley, J. W., & Goldberg, M. E. (2003). Neuronal activity in the lateral intraparietal area and spatial attention. Science (New York, N.Y.), 299(5603), 81–86.

Brown, R., Lau, H., & LeDoux, J. E. (2019). Understanding the higher-order approach to consciousness. Trends in Cognitive Sciences, 23(9), 754–768.

Buxbaum, L. J. (2006). On the right (and left) track: Twenty years of progress in studying hemispatial neglect. Cognitive Neuropsychology, 23(1), 184–201.

Buxbaum, L. J., & Coslett, H. B. (1994). Neglect of chimeric figures: two halves are better than a whole. Neuropsychologia, 32(3), 275–288.

Calvanio, R., Petrone, P. N., & Levine, D. N. (1987). Left visual spatial neglect is both environment-centered and body-centered. Neurology, 37(7), 1179–1183.

Caramazza, A., & Hillis, A. E. (1990). Levels of representation, co-ordinate frames, and unilateral neglect. Cognitive Neuropsychology, 7(5–6), 391–445.

Cohen, A., & Rafal, R. D. (1991). Attention and Feature Integration: Illusory Conjunctions in a Patient with a Parietal Lobe Lesion. Psychological Science, 2(2), 106–110.

Corbetta, M., Kincade, M. J., Lewis, C., Snyder, A. Z., & Sapir, A. (2005). Neural basis and recovery of spatial attention deficits in spatial neglect. Nature Neuroscience, 8(11), 1603–1610.

Corbetta, M., & Shulman, G. L. (2011). Spatial neglect and attention networks. Annual Review of Neuroscience, 34, 569–599.

Cowey, A. (1985). Aspects of cortical organization related to selective attention and selective impairments of visual perception: A tutorial review. Attention and Performance XI, 41–62.

Crick, F., & Koch, C. (1998). The problem of consciousness. In Consciousness and Emotion in Cognitive Science (pp. 63–69). Routledge.

De Renzi, E. (1982). Disorders of Space Exploration and Cognition. J. Wiley.

Deouell, L. Y. (2002). Pre-requisites for conscious awareness: clues from electrophysiological and behavioral studies of unilateral neglect patients. Consciousness and Cognition, 11(4), 546–567.

Desimone, R., & Duncan, J. (1995). Neural mechanisms of selective visual attention. Annual Review of Neuroscience, 18, 193–222.

Driver, J., & Mattingley, J. B. (1998). Parietal neglect and visual awareness. Nature Neuroscience, 1(1), 17–22.

Driver, J., & Vuilleumier, P. (2001). Perceptual awareness and its loss in unilateral neglect and extinction. Cognition, 79(1–2), 39–88.

D’Zmura, M. (1991). Color in visual search. Vision Research, 31(6), 951–966.

Esterman, M., McGlinchey-Berroth, R., Verfaellie, M., Grande, L., Kilduff, P., & Milberg, W. (2002). Aware and unaware perception in hemispatial neglect: evidence from a stem completion priming task. Cortex; a Journal Devoted to the Study of the Nervous System and Behavior, 38(2), 233–246.

Farne, A., Buxbaum, L. J., Ferraro, M., Frassinetti, F., Whyte, J., Veramonti, T., Angeli, V., Coslett, H. B., & Ladavas, E. (2004). Patterns of spontaneous recovery of neglect and associated disorders in acute right brain-damaged patients. Journal of Neurology, Neurosurgery, and Psychiatry, 75(10), 1401–1410.

Fellrath, J., & Ptak, R. (2015). The role of visual saliency for the allocation of attention: Evidence from spatial neglect and hemianopia. Neuropsychologia, 73, 70–81.

Ferster, D., & Miller, K. D. (2000). Neural mechanisms of orientation selectivity in the visual cortex. Annual Review of Neuroscience, 23(1), 441–471.

Friedman-Hill, Friedman-Hill, Robertson, L., & Treisman, A. (1995). Parietal contributions to visual feature binding: evidence from a patient with bilateral lesions. In Science (Vol. 269, Issue 5225, pp. 853–855). https://doi.org/10.1126/science.7638604

Grill-Spector, K., Kushnir, T., Hendler, T., Edelman, S., Itzchak, Y., & Malach, R. (1998). A sequence of object-processing stages revealed by fMRI in the human occipital lobe. Human Brain Mapping, 6(4), 316–328.

Halligan, P. W., Burn, J. P., Marshall, J. C., & Wade, D. T. (1992). Visuo-spatial neglect: qualitative differences and laterality of cerebral lesion. Journal of Neurology, Neurosurgery, and Psychiatry, 55(11), 1060–1068.

Hillis, A. E., & Caramazza, A. (1995). A framework for interpreting distinct patterns of hemispatial neglect. Neurocase, 1(3), 189–207.

Hubel, D. H., & Wiesel, T. N. (1962). Receptive fields, binocular interaction and functional architecture in the cat’s visual cortex. The Journal of Physiology, 160, 106–154.

Hubel, D. H., & Wiesel, T. N. (1968). Receptive fields and functional architecture of monkey striate cortex. The Journal of Physiology, 195(1), 215–243.

Humphreys, G. W., Cinel, C., Wolfe, J., Olson, A., & Klempen, N. (2000). Fractionating the binding process: neuropsychological evidence distinguishing binding of form from binding of surface features. Vision Research, 40(10–12), 1569–1596.

Hutchinson, B. T. (2019). Toward a theory of consciousness: A review of the neural correlates of inattentional blindness. Neuroscience and Biobehavioral Reviews, 104, 87–99.

Kahneman, D., Treisman, A., & Gibbs, B. J. (1992). The reviewing of object files: object-specific integration of information. Cognitive Psychology, 24(2), 175–219.

Kinsbourne, M. (1987). Mechanisms of Unilateral Neglect. In M. Jeannerod (Ed.), Advances in Psychology (Vol. 45, pp. 69–86). North-Holland.

Kinsbourne, M., & Warrington, E. K. (1962). A variety of reading disability associated with right hemisphere lesions. Journal of Neurology, Neurosurgery, and Psychiatry, 25, 339– 344.

Kourtzi, Z., & Kanwisher, N. (2001). Representation of perceived object shape by the human lateral occipital complex. Science, 293(5534), 1506–1509.

Kristjánsson, A., Vuilleumier, P., Malhotra, P., Husain, M., & Driver, J. (2005). Priming of color and position during visual search in unilateral spatial neglect. Journal of Cognitive Neuroscience, 17(6), 859–873.

Làdavas, E., Carletti, M., & Gori, G. (1994). Automatic and voluntary orienting of attention in patients with visual neglect: horizontal and vertical dimensions. Neuropsychologia, 32(10), 1195–1208.

Làdavas, E., Paladini, R., & Cubelli, R. (1993). Implicit associative priming in a patient with left visual neglect. In Neuropsychologia (Vol. 31, Issue 12, pp. 1307–1320). https://doi.org/10.1016/0028-3932(93)90100-e

Livingstone, M., & Hubel, D. (1988). Segregation of form, color, movement, and depth: anatomy, physiology, and perception. Science, 240(4853), 740–749.

Lundqvist, D., Flykt, A., & Öhman, A. (1998). The Karolinska directed emotional faces (KDEF). CD ROM from Department of Clinical Neuroscience, Psychology Section, Karolinska Institutet, 91(630), 2–2.

Lunven, M., & Bartolomeo, P. (2017). Attention and spatial cognition: Neural and anatomical substrates of visual neglect. Annals of Physical and Rehabilitation Medicine, 60(3), 124– 129.

Mack, A., & Rock, I. (2000). Inattentional Blindness. Bradford Books.

McGlinchey-Berroth, R. (1997). Visual information processing in hemispatial neglect. In Trends in Cognitive Sciences (Vol. 1, Issue 3, pp. 91–97). https://doi.org/10.1016/s1364-6613(97)89054-7

Medina, J., Kannan, V., Pawlak, M. A., Kleinman, J. T., Newhart, M., Davis, C., Heidler-Gary, J. E., Herskovits, E. H., & Hillis, A. E. (2009). Neural substrates of visuospatial processing in distinct reference frames: evidence from unilateral spatial neglect. Journal of Cognitive Neuroscience, 21(11), 2073–2084.

Palmer, S., & Rock, I. (1994). Rethinking perceptual organization: The role of uniform connectedness. Psychonomic Bulletin & Review, 1(1), 29–55.

Petrides, M., & Pandya, D. N. (2002). Association pathways of the prefrontal cortex and functional observations. In Principles of Frontal Lobe Function (pp. 31–50). Oxford University Press.

Pitzalis, S., Spinelli, D., & Zoccolotti, P. (1997). Vertical neglect: Behavioral and electrophysiological data. Cortex; a Journal Devoted to the Study of the Nervous System and Behavior, 33(4), 679–688.

Rafal, R. D. (1994). Neglect. Current Opinion in Neurobiology, 4(2), 231–236.

Rees, G. (2001). Neuroimaging of visual awareness in patients and normal subjects. Current Opinion in Neurobiology, 11(2), 150–156.

Rees, G., Kreiman, G., & Koch, C. (2002). Neural correlates of consciousness in humans. Nature Reviews. Neuroscience, 3(4), 261–270.

Rensink, R. A., O’Regan, J. K., & Clark, J. J. (1997). To see or not to see: The need for attention to perceive changes in scenes. Psychological Science, 8(5), 368–373.

Rizzolatti, & Gallese. (1988). Mechanisms and theories of spatial neglect. Handbook of Neuropsychology, 1, 223–246.

Ro, T., & Rafal, R. D. (1996). Perception of geometric illusions in hemispatial neglect. Neuropsychologia, 34(10), 973–978.

Robertson, L. C. (2003). Binding, spatial attention and perceptual awareness. Nature Reviews. Neuroscience, 4(2), 93–102.

Roelfsema, P. R. (2006). Cortical algorithms for perceptual grouping. Annual Review of Neuroscience, 29(1), 203–227.

Schubert, T., & McCloskey, M. (2013). Prelexical representations and processes in reading: evidence from acquired dyslexia. Cognitive Neuropsychology, 30(6), 360–395.

Snodgrass, J. G., & Vanderwart, M. (1980). A standardized set of 260 pictures: norms for name agreement, image agreement, familiarity, and visual complexity. Journal of Experimental Psychology. Human Learning and Memory, 6(2), 174–215.

Subbiah, I., & Caramazza, A. (2000). Stimulus-centered neglect in reading and object recognition. Neurocase, 6(1), 13–31.

Thiebaut de Schotten, M., Urbanski, M., Duffau, H., Volle, E., Lévy, R., Dubois, B., & Bartolomeo, P. (2005). Direct evidence for a parietal-frontal pathway subserving spatial awareness in humans. Science (New York, N.Y.), 309(5744), 2226–2228.

Treccani, B., Cubelli, R., Sellaro, R., Umiltà, C., & Della Sala, S. (2012). Dissociation between awareness and spatial coding: evidence from unilateral neglect. Journal of Cognitive Neuroscience, 24(4), 854–867.

Treisman, A. M. (1986). Handbook of human perception and performance (K. R. Boff, L. Kaufmann, & J. P. Thomas, Eds.; Vol. 2, pp. 37–38). John Wiley & Son.

Treisman, A. M. (2006). How the deployment of attention determines what we see. Visual Cognition, 14(4–8), 411–443.

Treisman, A. M., & Gelade, G. (1980). A feature-integration theory of attention. Cognitive Psychology, 12(1), 97–136.

Treisman, A. M., & Souther, J. (1985). Search asymmetry: a diagnostic for preattentive processing of separable features. Journal of Experimental Psychology. General, 114(3), 285–310.

Vallar, G., & Calzolari, E. (2018). Unilateral spatial neglect after posterior parietal damage. Handbook of Clinical Neurology, 151, 287–312.

Vallar, G., Daini, R., & Antonucci, G. (2000). Processing of illusion of length in spatial hemineglect: a study of line bisection. Neuropsychologia, 38(7), 1087–1097.

Vallar, G., & Maravita, A. (2003). Handbook of neuroscience for the behavioral sciences (G. G. Berntson & J. T. Cacioppo, Eds.; Vol. 1, pp. 322–336). Pierre Bonnier’s.

Vallar, G., & Perani, D. (1987). The Anatomy of Spatial Neglect in Humans. In M. Jeannerod (Ed.), Advances in Psychology (Vol. 45, pp. 235–258). North-Holland.

Van Vleet, T. M., & Robertson, L. C. (2009). Implicit representation and explicit detection of features in patients with hemispatial neglect. Brain: A Journal of Neurology, 132(Pt 7), 1889–1897.

Vandenbroucke, A. R. E., Fahrenfort, J. J., Sligte, I. G., & Lamme, V. A. F. (2014). Seeing without knowing: neural signatures of perceptual inference in the absence of report. Journal of Cognitive Neuroscience, 26(5), 955–969.

Vannuscorps, G., Galaburda, A., & Caramazza, A. (2021). Shape-centered representations of bounded regions of space mediate the perception of objects. Cognitive Neuropsychology, 1–50.

Verdon, V., Schwartz, S., Lovblad, K.-O., Hauert, C.-A., & Vuilleumier, P. (2010). Neuroanatomy of hemispatial neglect and its functional components: a study using voxel-based lesion-symptom mapping. Brain: A Journal of Neurology, 133(Pt 3), 880–894.

Vuilleumier, P., Sagiv, N., Hazeltine, E., Poldrack, R. A., Swick, D., Rafal, R. D., & Gabrieli, J. D. (2001). Neural fate of seen and unseen faces in visuospatial neglect: a combined event-related functional MRI and event-related potential study. Proceedings of the National Academy of Sciences of the United States of America, 98(6), 3495–3500.

Wolfe, J. M., & Bennett, S. C. (1997). Preattentive object files: shapeless bundles of basic features. Vision Research, 37(1), 25–43.

Wolfe, J. M., Friedman-Hill, S. R., Stewart, M. I., & O’Connell, K. M. (1992). The role of categorization in visual search for orientation. Journal of Experimental Psychology. Human Perception and Performance, 18(1), 34–49.

Wolfe, J. M., & Horowitz, T. S. (2004). What attributes guide the deployment of visual attention and how do they do it? Nature Reviews. Neuroscience, 5(6), 495–501.

Xu, Y. (2017). Reevaluating the Sensory Account of Visual Working Memory Storage. Trends in Cognitive Sciences, 21(10), 794–815.

Xu, Y. (2018). The Posterior Parietal Cortex in Adaptive Visual Processing. Trends in Neurosciences, 41(11), 806–822.

Young, A. W., Hellawell, D. J., & Welch, J. (1992). NEGLECT AND VISUAL RECOGNITION. Brain: A Journal of Neurology, 115(1), 51–71.

Young, A. W., Newcombe, F., & Ellis, A. W. (1991). Different impairments contribute to neglect dyslexia. Cognitive Neuropsychology, 8(3–4), 177–191.

Zeki, S. (1993). A vision of the brain. Blackwell scientific publications.

Zeki, S., Watson, J. D., Lueck, C. J., Friston, K. J., Kennard, C., & Frackowiak, R. S. (1991). A direct demonstration of functional specialization in human visual cortex. The Journal of Neuroscience: The Official Journal of the Society for Neuroscience, 11(3), 641–649.

