## Supplementary material for "sPreserved recognition of basic visual features despite lack of awareness of shape: evidence from a case of neglect": Supplemantary Text

**Supplemental Information**

**Appendix A – General neuropsychological screening**

XX's speech production and comprehension are normal. He performed at ceiling (60/60 correct) in confrontation naming of line drawings of objects on the Boston Naming Task and his description of the Cookie Theft Picture from the Boston Diagnostic Aphasia Examination (an image depicting a family in a kitchen) was appropriate, elaborate, and detailed (Goodglass & Kaplan, 1983). His verbal short-term memory was also within the normal range with a forward digit span of 8 and a backward digit span of 4 (for age and education matched norms, see Shirk et al., 2011).

His lexical processing for written words was systematically assessed with word lists compiled from the Johns Hopkins University Dysgraphia and Dyslexia Batteries (Goodman & Caramazza, 1985) and custom word lists. When administered an 80-item list of 4- or 5-letter words mixed with pronounceable nonwords and asked to name them in response to oral spelling, he performed at ceiling (77/80 correct) suggesting intact *reading* mechanisms when visual input is bypassed (Schubert & McCloskey, 2015). His written spelling to dictation was assessed with a 48-item word list (4 to 7 letters); he made no errors, suggesting that he has intact knowledge of lexical orthographic representations.

XX also did not present any difficulties in basic visual perception aside from his neglect. Tested with several subtests of the Birmingham Object Recognition Battery (BORB, Riddoch & Humphreys, 1993), he performed within the normal range (except for neglect symptoms). The BORB overlapping figures subtest revealed intact figure-ground separation and grouping: When presented with line drawings of everyday objects presented individually (40/40 correct), in pairs (20 pairs, 40/40 objects correct), or overlapping (20 overlapping pairs, 40/40 objects correct), he correctly named all items. He was also within the normal range (23/25 correct) at perceiving and identifying objects in three dimensions (BORB foreshortened view subtest).

**Appendix B – Neglect screening**

Screening of neglect was carried out using standard screening batteries (unlimited duration) as well as custom experiments (duration limited). Despite relatively good performance on some neglect screening tests with unlimited duration, XX’s performance was clearly abnormal. In line cancellation and letter cancellation (with items randomly dispersed across the page), he successfully canceled all items on both sides of the page. However, when the letters were presented as two columns, he missed 2/20 letters in the left column. In the bells test (various small line drawings are randomly dispersed on a page and the patient is asked to cross the bells among the distractors), he missed 6/17 bells on the left. In these tasks, he mostly started searching for items on the right and moved to the left after completing the right side. In line bisection, he showed a rightward bias in 3 of 4 trials (8%, 16%, 3% of the line). In direct copying of a single flower, he missed the petals on the left (see Figure S1). Custom computerized tasks with limited exposure durations allowed us to gain a more detailed understanding of XX’s hemispatial neglect.

**Appendix C - Determining the affected reference frame**

XX had difficulty in perceiving visual information on the left half of words, objects, and faces in a variety of tasks with stimuli presented centrally. However, when stimuli are presented centrally and in their canonical orientation, all reference frames are aligned, making it impossible to distinguish the level affected by neglect. To determine the affected reference frame, we ran a series of experiments manipulating the position and orientation of stimuli (Experiments 4a-4e, see Supplemental Methods for details).

One main distinction of reference frames is between retinocentric and non-retinocentric. A simple test of whether the deficit is retinocentric is to present stimuli to the right or left of fixation (in the right or left visual field). With a retinocentric deficit, stimuli falling in the left visual field will be neglected but stimuli in the right visual field will be processed normally. However, if the deficit occurs at a non-retinocentric reference frame, left-sided errors should persist for both right and left visual fields. To test for a retinocentric deficit, we first used a simple visual shape detection paradigm and presented black circles to the left and/or right of fixation (see Experiment 4a). XX had no difficulty in detecting the circles in the left visual field.

Next, we presented words to the left or right of fixation to test his word reading across the two visual fields (see Experiment 4b, SI Figure S3). He was more likely to make a word reading error when the word was presented on the left (left: 53% error, right: 26% error, *X^2^* = 6.77, *p* = .009). However, regardless of the word’s position, he made similar left-sided letter substitution errors (e.g., reading ‘*calf*’ as ‘*self*, see SI Figure S3). That is, even when the entirety of the word fell on the right visual field, he continued to make errors on the left half of the word.

As a final test of a retinocentric deficit, we displayed two stimuli bilaterally to the left or right of fixation and asked him to report if they are the same (Experiment 4c). When the right halves of the stimuli were identical, he was at chance in determining whether they are the same or different (see SI Figure S4; same/different task: 59% correct for identical pairs, at chance, *p* = .63; 61% correct for pairs that differed on their left sites, at chance, *p* = .48; 85% correct for pairs that differed on both sides, above chance, *p* = .003). Overall, he responded as if he did not have access to the left halves of stimuli in both left and right visual fields. These patterns suggest a non-retinocentric deficit: stimulus-centered or object/word-centered.

To assess whether his deficit is at stimulus- or object-centered levels of processing, we presented stimuli in their non-canonical orientation; object-centered left neglect affects the left side of a canonically-oriented representation of a visual stimulus – e.g., the beginning of words in English irrespective of position or orientation. For instance, when a word is rotated 180 degrees, the beginning of the word would fall on the right side of the stimulus, and an individual with object-centered left neglect would make errors at the beginning of a word. XX, on the other hand, tended to make letter substitution errors at the end of the word, falling to the left side of the stimulus (Experiment 4d, see SI Figure S5, e.g., reading *rush* as *runs*). This pattern suggests that XX’s deficit does not concern the object/word-centered level of representation.

We also tested the effect of stimulus orientation by using objects and chimeric figures tilted 45 degrees counterclockwise (Experiment 4e, see SI Figure S6). Tilt would not affect performance at an object-centered reference frame, in which objects are normalized to a standard (upright) position. However, tilting the figure 45 degrees changes the horizontal spatial extent of the figure affecting the representation at the stimulus-based level, shifting some of the information on the left side to the non-neglected right half. XX’s performance in the object decision task improved when the chimeric figures were tilted: he correctly recognized 72% of chimeric figures as nonreal (above chance, *p* = .048), a significant increase from his 6% performance on the same stimuli presented upright (compare Figure 1A and SI Figure S6).

Overall, we found XX’s error patterns to be affected by orientation (Experiments 4d,e) and unaffected by position in the visual field (Experiment 4a-c), leading us to conclude that his neglect operates at a stimulus-centered reference frame. At this level, the coordinate frame is aligned to the stimulus defined by its spatial extent, rather than positions in the visual field (e.g., Hillis & Caramazza, 1995; Ellis et al., 1987; Subbiah & Caramazza, 2000). Stimulus-centered neglect is expected to affect the left halves of individually segmented objects, and the problem in processing the left half persists even when the entire stimulus falls to the right visual field.

**Supplemental Methods**

**Experiment 4a.** Black filled circles (90 pixels radius) were presented bilaterally and on the left or right of a fixation cross. The stimuli stayed on the screen for 500, 1000, or 2000 milliseconds (a total of 60 items, 20 items for each duration, equally distributed across the three conditions). XX was asked to maintain fixation and report whether he saw one, two, or zero circles. Regardless of the number of stimuli, position in the visual field, or exposure duration, he detected all items, including left circles shown in a bilateral display, indicating that he does not have extinction, which commonly co-occurs with neglect (Umarova et al., 2011). Furthermore, that he could detect stimuli that are presented on the left visual field suggests that he does not have a retinocentric deficit.

**Experiment 4b.** One hundred twenty 4-letter words with different numbers of neighbors or cohort words were presented in the left or right of the visual field (60 on the left, 60 on the right), in a 50-point size, set in lowercase Courier font and against a white background. XX was asked to maintain fixation and the words were presented for 800 milliseconds to mitigate the effect of seeing the words in the periphery. The words were presented centrally in their respective half of the visual field. XX’s task was to maintain fixation and read the words (valid trials: 49/60 left side, 54/60 right side).

**Experiment 4c.** Chimeric figures or objects from Experiment 1b were presented bilaterally on the left or right of a fixation cross (see SI Figure S4, 60 trials in total: 20 pairs that differed on their left sides, 20 identical pairs, and 20 pairs that differed on both sides). XX maintained fixation during the task and was asked to report if the stimuli were the same or different (valid trials: 20/20 both-halves-different pairs, 17/20 identical pairs, 18/20 left-halves-different pairs).

**Experiment 4d.** A list of 40 4-letter words mixed with pronounceable nonwords were shown at fixation in upright position for 500 milliseconds, or rotated 180 degrees, for 800 milliseconds (to mitigate the effect of seeing letters in a noncanonical orientation, see SI Figure S5; valid trials: 39/40 upright, 37/40 rotated). The words were presented in a 50-point size, set in lowercase Courier font and against a white background.

**Experiment 4e.** Chimeric figures or real objects from Experiment 1a were presented tilted 45 degrees counterclockwise (see SI Figure S6, 40 trials in total: 20 unique stimuli (10 real) and their reflections). XX’s task was to report whether the stimuli were real (valid trials: 18/20 for real objects, 18/20 for chimeric figures).

**Supplemental Video Captions**

Supplemental videos are deposited at the Open Science Framework (<https://osf.io/nbzwu/>).

**Video S1: Object decision of a black and white chimeric figure.** In this video, XX is a presented with a chimeric figure that is half a piano (left) and half a sailboat (right). He described this figure as a “*real sailboat*”.

**Video S2: Word reading.** Example word reading performance of XX when a word is presented centrally. In this video, XX reads the word *stim* as “grim”.

**Video S3: Object decision and color naming of a bicolor chimeric figure.** In this video, XX is presented with a bicolor chimeric figure that is half a swan (left, red) and half a truck (right, blue). He describes this figure as “*a truck that is red on the left and blue on the right*”.

**Video S4: Colored word reading.** In this video, XX is presented with the word *cosy* where the letters ‘c’ and ‘o’ are red, and ‘s’ and ‘y’ are blue. He reads the word as “*easy*” but can accurately report the color on both sides.

**Supplemental Figures**

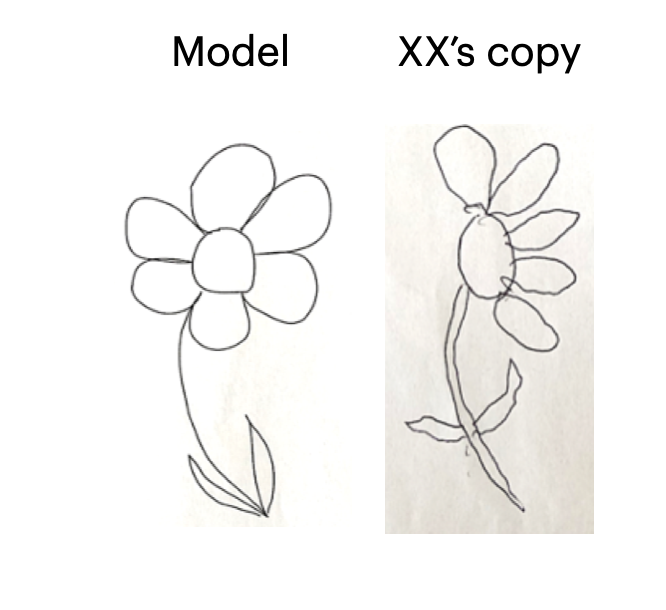

**Figure S1.** Sample response in a direct-copying task of a single flower. XX had unlimited duration to complete this task.

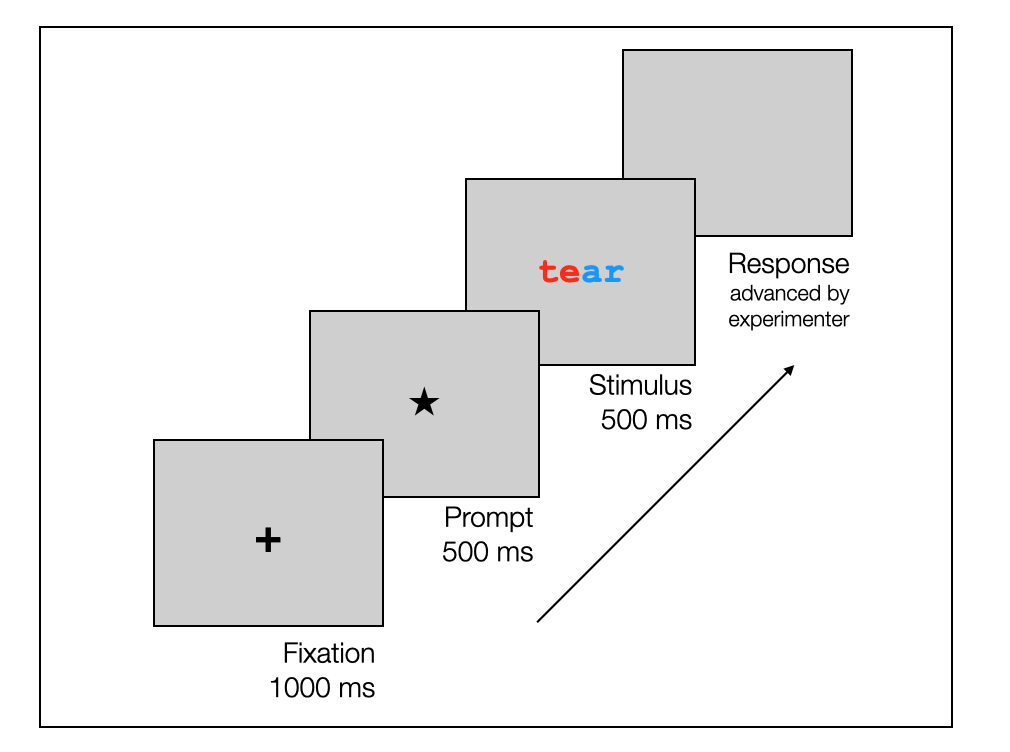

**Figure S2.** Illustration of the display sequence used in the custom computerized tasks. Every trial started with a central fixation cross (1000 ms), followed by a prompt (square or a star, 500 ms). The stimulus followed the square/star prompt and stayed on the screen for 500 milliseconds. After the stimulus, a blank screen appeared where XX provided his response. The blank screen stayed on the screen until the response period was over and was advanced by the experimenter. Only those trials where XX correctly reported the square or star prompt were included as valid trials for analyses. The trials where he did not report the prompt were interpreted as inattention to task, or disruption of fixation right before the stimulus, and excluded from the analyses.

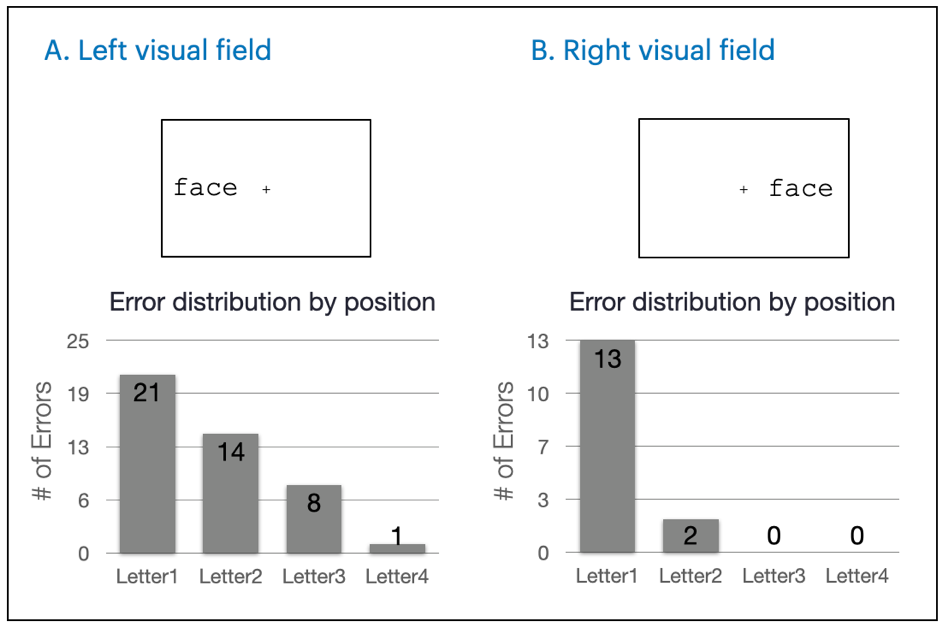

**Figure S3.** Distribution of letter substitution errors as a function of position in the visual field and letter position. He was more likely to make a word reading error when the word was presented on the left (left: 53% error, right: 26% error, *X^2^* = 6.77, *p* = .009). However, he tended to make left-sided letter substitution errors in both cases, and his erroneous responses generally preserved the word length (left: 93%, right: 93% of erroneous responses). For ease of comparison across different letter positions, only the errors that matched the target word length are included in the graph.

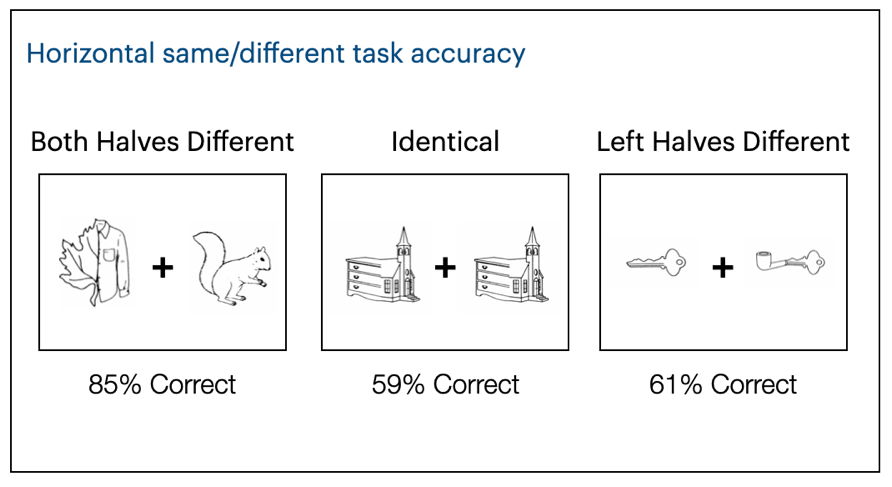

**Figure S4.**Same-different task performance for objects or chimeric figures presented side by side. XX performed at chance when the two stimuli were identical or only different on the left, but he was able to distinguish the images when the difference was marked on both sides.

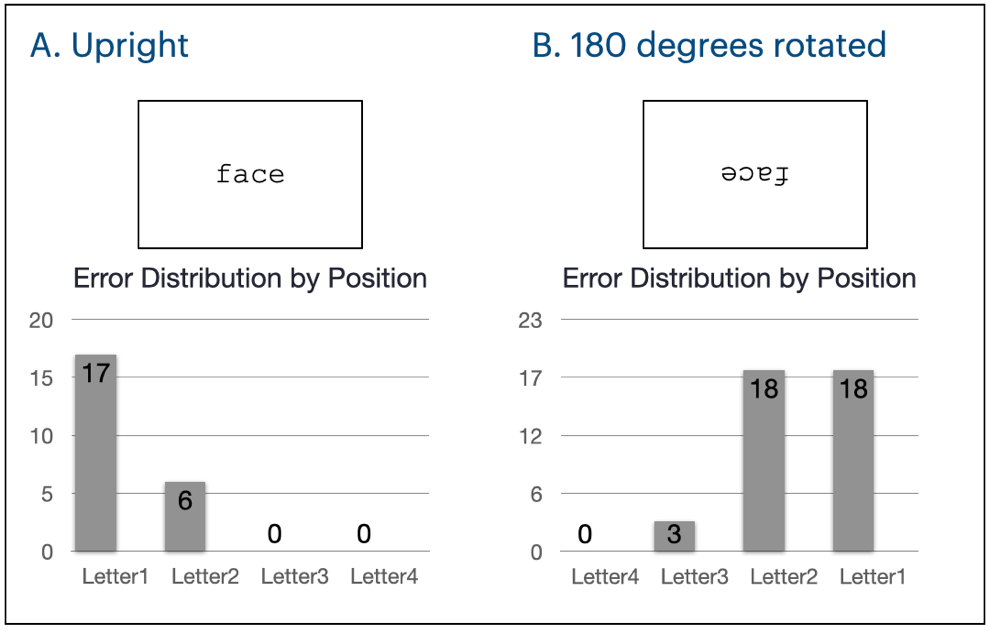

**Figure S5.** Distribution of lexical substitution errors of by letter position for words presented upright or 180 degrees rotated. For ease of comparison across different letter positions, only the errors that matched the target word length are included in the graph (upright: 100%, inverted: 92% of erroneous responses). For upright words, he tended to make letter substitution errors at the beginning of the word, falling to the left side (51% error, e.g., reading *sank* as *sink*). In the rotated condition, he made letter substitution errors at the end of the word, falling to the left side (68% error, e.g., reading *sank* as *safe*). This pattern indicates that XX’s deficit does not concern a word/object-centered frame of reference.

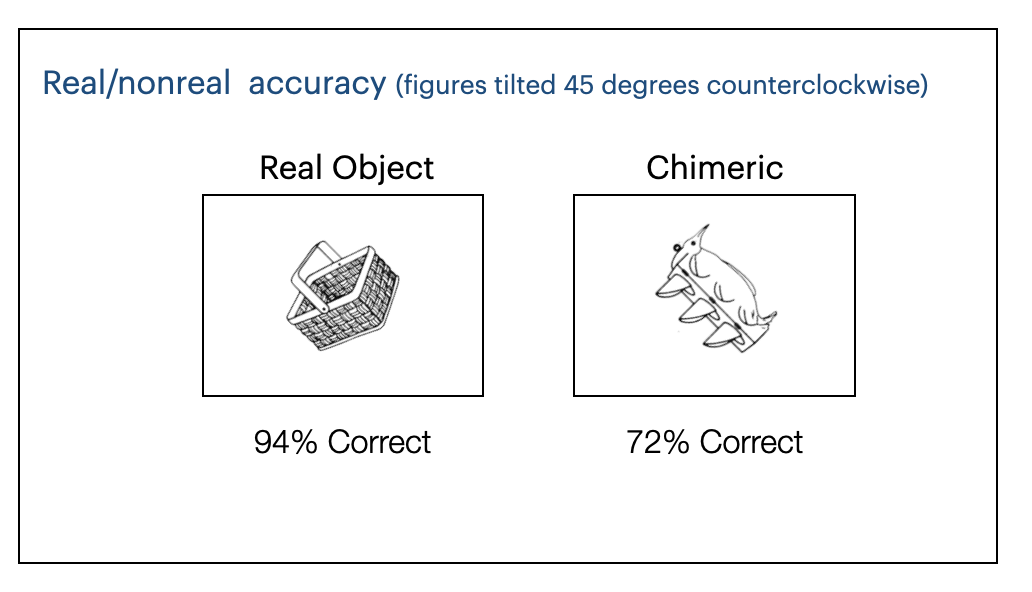

**Figure S6.** Real/nonreal object decision performance for chimeric figures or objects tilted 45 degrees counterclockwise. XX’s performance in the real/nonreal object decision task improved when the objects were tilted: he correctly recognized 72% of chimeric figures as nonreal, an increase from his 6% performance on the same stimuli presented upright (compare to Figure 1A).

**Supplementary References**

Goodglass, H., & Kaplan, E. (1983). *Boston diagnostic aphasia examination booklet*. Lea & Febiger.

Goodman, R. A., & Caramazza, A. (1985). The Johns Hopkins university dysgraphia battery. *Baltimore, MD: Johns Hopkins University*.

Riddoch, J. M., & Humphreys, G. W. (1993). *BORB: Birmingham Object Recognition Battery*. Taylor & Francis Group.

Schubert, T., & McCloskey, M. (2015). Recognition of oral spelling is diagnostic of the central reading processes. *Cognitive Neuropsychology*, *32*(2), 80–88.

Shirk, S. D., Mitchell, M. B., Shaughnessy, L. W., Sherman, J. C., Locascio, J. J., Weintraub, S., & Atri, A. (2011). A web-based normative calculator for the uniform data set (UDS) neuropsychological test battery. *Alzheimer’s Research & Therapy*, *3*(6), 32.

Umarova, R. M., Saur, D., Kaller, C. P., Vry, M.-S., Glauche, V., Mader, I., Hennig, J., & Weiller, C. (2011). Acute visual neglect and extinction: distinct functional state of the visuospatial attention system. *Brain: A Journal of Neurology*, *134*(Pt 11), 3310–3325.
